# HIV-1 Nef uses a conserved pocket to recruit the N-terminal cytoplasmic tail of Serinc3

**DOI:** 10.1101/2025.04.12.648528

**Authors:** Mohammad Karimian Shamsabadi, Charlotte Stoneham, Amalia De-leon, Tony Fares, John Guatelli, Xiaofei Jia

## Abstract

Human transmembrane proteins Serinc3 and Serinc5 are antiviral restriction factors that inhibit HIV-1 infectivity. In the absence of viral antagonism, Serinc3 and Serinc5 incorporate into the envelopes of nascent virions and inhibit the fusion of virions to the target cells. The HIV-1 virus counteracts the restriction of Serinc3 by downregulating it from the cell surface and thus excluding it from budding virions. This is orchestrated by the viral accessory protein Nef and involves hijacking of the clathrin adaptor protein complex 2 (AP2)-dependent endocytosis. The mechanistic details of Nef-mediated Serinc3 downregulation, however, have been enigmatic. In this work, we investigated and revealed the molecular determinants of Serinc3 modulation by Nef. Our results show that Nef recruits Serinc3 by binding to its N-terminal cytosolic tail. Furthermore, Nef residues important for Serinc3-binding *in vitro,* and for the exclusion of Serinc3 from virions, overlap with those required for Nef-mediated CD4 downregulation, suggesting great mechanistic similarities between the two functions of Nef. In addition to shedding light to the mechanism of Serinc3 antagonism, our work also highlights the conserved substrate-binding pocket of Nef as a molecular hotspot for inhibitor development and antiretroviral drug discovery.

## Introduction

Serinc3 and Serinc5 proteins are members of the Serinc family, which additionally includes Serinc1, 2, and 4 (1). Serinc proteins are multipass transmembrane proteins whose normal cellular functions are incompletely understood (2). In 2015, Serinc3 and Serinc5 were identified, through two independent studies, as restriction factors of HIV-1 infection (3, 4). In the absence of viral antagonism, Serinc3 and Serinc5 are incorporated into the envelopes of the budding virions leading to reduction of the virions’ infectivity (3, 4). Specifically, their presence inhibits the fusion between the lipid membrane of the incoming virion and that of the target cell (3–6). Not all strains of HIV-1 are subjected to restriction by Serinc3 and Serinc5; whether, or to what extent, the restriction occurs depends on the viral Env protein (5, 7). For the sensitive strains, the conformation of Env was found to be altered by the virion-incorporated Serinc3/5 (5, 8–10), although this effect might not be caused by a direct Env-Serinc interaction (5). While Serinc5 expression does not change the lipid composition of the viral or host cell membranes (11), Serinc3 and Serinc5 are lipid transporters that reduce the asymmetry of the lipid membrane (12). Such lipid flipping activities of Serinc3/5 were shown to correlate with their restriction activities as well as with conformational changes of Env, providing a clue to the mechanistic basis of Serinc3/5’s antiviral activities (12).

The antiviral activities of Serinc3 and Serinc5 are antagonized by the HIV-1 accessory protein Nef (3, 4). Nef, a multifunctional peripheral membrane protein, is expressed early in the viral replication cycle and plays a key role in viral pathogenesis (13). It modulates T cell activation through direct binding with cellular kinases, which presumably favors HIV-1 replication (14, 15). In addition, Nef downregulates immune-receptor molecules from the cell surface, enabling infected cells to evade both adaptive and innate immunity (13, 16). The best characterized Nef activities are downregulation of cell-surface CD4, which enables infected cell to evade antibody-dependent cellular cytotoxicity (ADCC) (17–20), and cell-surface downregulation of MHC-I, which hides infected cells from immune surveillance by cytotoxic T lymphoctytes (CTLs) (21). Nef executes the downregulation of surface CD4 by hijacking the clathrin AP2-dependent endocytosis (22, 23), while downregulation of MHC-I takes place through Nef-mediated hijacking of clathrin AP1-dependent membrane trafficking (24–27).

Counteraction of Serinc3 and Serinc5 by Nef, like that of CD4, also occurs at the plasma membrane and similarly involves hijacking of clathrin AP2-dependent endocytosis (4, 28, 29). AP2, the mediator of clathrin coats formation at the plasma membrane, is a tetrameric complex containing two large subunits (α and β2), one medium subunit (μ2), and one small subunit (σ2). AP2 recruits membrane cargos through binding to the sorting motifs located within the cytoplasmic domains of the cargo proteins. Two classes of motifs are commonly recognized by AP2 (as well as by other APs): the tyrosine-based motifs denoted as YxxΦ (Φ: a large hydrophobic residue: x: any amino acid) and the acidic dileucine motifs denoted as (E/D)xxxL(L/I) (30). Nef hijacks AP2 partly through mimicking the acidic dileucine sorting motif: as revealed by high-resolution structures the ExxxLL sequence located within the long and flexible C-terminal loop of Nef binds into a site on the σ2 and α subunits of AP2 dedicated to acidic dileucine motif-binding (Figure 1AB) (31, 32). The rest of Nef’s C-terminal loop also makes substantial interactions with the σ2 subunit of AP2 (31, 32). The extensive binding between Nef’s C-terminal loop and AP2, which has been observed with or without CD4 bound to Nef (31, 32), is the foundation for the hijacking of the clathrin-mediated endocytosis by Nef.

**Figure 1.**
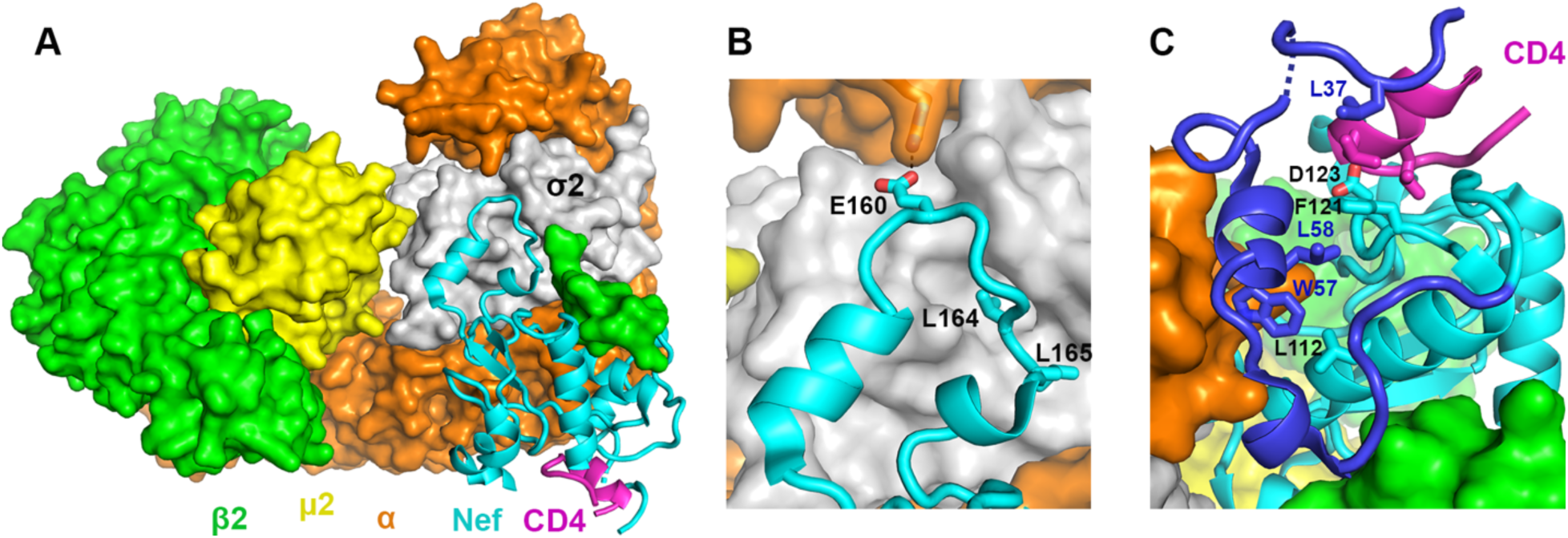
Mechanistic insights on Nef-mediated clathrin AP2-dependent CD4 downregulation gained by our previous crystal structure (PDB ID: 6URI) (32). A) Nef “connects” CD4 to the heterotetrameric clathrin AP2 complex. B) The C-terminal loop of Nef binds extensively to the σ2 and α subunits of AP2 partly by mimicking the acidic dileucine motif. C) Nef’s N-terminal loop (dark blue) adopts a unique conformation in the complex. Part of the loop forms a wall of the CD4-binding pocket with Leu37 making direct contacts with CD4. Phe121 and Asp123 on the core domain of Nef also interact with CD4. Nef residues Trp57 and Leu58 mediate the docking of the short helix into the hydrophobic pocket formed at the Nef–α interface.

While engaging AP2 through its C-terminal loop, Nef uses a different pocket to bind the cytoplasmic domain of CD4 (Figure 1A), as revealed by our previous structure (32). Thus, Nef functions as a “connector” between AP2 and CD4. CD4-binding involves residues both from Nef’s rigid core (e.g., Phe121 and Asp123) and from its N-terminal loop (Leu37) (Figure 1C) (32). Formation of this CD4-binding pocket depends on the Nef’s N-terminal loop adopting a unique conformation (Figure 1C). Here, two residues on the N-terminal loop—Trp57 and Leu58—dock into a shallow hydrophobic pocket on the Nef core making direct contact with Leu112 and Phe121 (32). Notably, mutations of these Nef residues, which are involved either in CD4-binding or in stabilizing the unique Nef conformation, indeed disrupted Nef-mediated surface downregulation of CD4 (22, 23, 32–36).

Downregulation of Serinc3 and Serinc5, which are similarly achieved through Nef-mediated hijacking of clathrin AP2-dependent endocytosis, prevents the incorporation of these innate restriction factors into virions and thus preserves virion-infectivity. Some mechanistic insights have so far been gained on the Serinc5-Nef interaction (2). An intracellular loop of Serinc5, namely intracellular loop 4 or ICL4, is targeted by Nef for binding and downregulation (37).

Within ICL4, residues Ile350 and Leu353 are required (37). On the Nef side, while several determinants were found to be shared between the Serinc5 downregulation and the CD4 downregulation (11, 38–41), some Nef residues were shown to be required for one downregulation but not the other, indicating distinction between the determinants in Nef for these two functions (42–45). When it comes to the downregulation of Serinc3, however, the mechanism is largely enigmatic. It is not known which segment of Serinc3 is targeted by Nef, which residues or surface(s) of Nef are involved in recruiting Serinc3, or which conformation of Nef is responsible for this function. To fill this knowledge gap, here we investigated and revealed the molecular determinants of Nef-mediated Serinc3 antagonism.

## MATERIALS AND METHODS

### Accession ID

Proteins involved in this work, and their associated UniProt IDs, are listed below: HIV-1 Nef (Q90VU7); Serinc3 (Q13530); AP2 α subunit (P18484); AP2 σ2 subunit (P53680); AP2 β2 subunit (P63010); AP2 μ2 subunit (P96CW1).

### Fusion construct design & protein expression and purification

Genes encoding the Serinc3 intracellular loops (NTL: 1-39; ICL1: 120-131; ICL2: 181-204; ICL3: 259-267; ICL4: 354-404) were each cloned into the pMAT9 vector. *E. Coli* cells transformed with the plasmid were grown at 37°C till OD_600_ reached 0.8. Protein expression was then induced with 0.1 mM IPTG and continued at 16°C overnight. Harvested cells were lysed using sonication. The protein of interest was purified sequentially through an MBP affinity column, a HiTrap Q anion exchange column, and finally a Superdex 200 size exclusion column.

The construct of MBP-Nef was created by cloning the gene encoding 26-206 of NL4.3 Nef into a pMAT9 vector. Protein expression and purification follow the same steps as above. Nef protein used in the GST pulldown assay was prepared by cleaving off the MBP tag from the MBP-Nef fusion protein, followed by a final purification through the HiTrap Q anion exchange column.

For the GST pulldown assay, the NTL-β2 fusion was constructed by fusing NTL to the N terminus of β2 subunit (1–591) of AP2 *via* a flexible linker of 10 amino acids. Genes encoding the above NTL-β2 fusion and the N-terminal domain of the µ2 subunit (1–135) of AP2 were cloned, respectively, into the two multiple cloning sites of the pETDuet expression vector. Production of the GST-tagged tetrameric NTL-AP2 complex was done by co-expressing NTL-β2 and µ2-NTD together with α(1–621)-GST and σ2 subunits of AP2 (encoded by genes carried in a pCDFDuet vector as described before (32)). *E. coli* cells transformed with these plasmids were grown at 37°C till OD_600_ reached 0.8. Protein expression was then induced with 0.1 mM IPTG and continued at 22°C overnight. Cells were then lysed using sonication. The protein of interest was purified using a Ni-NTA affinity column, followed by a GST affinity column. Purified protein was then buffer exchanged to a binding buffer free of glutathione (50 mM Tris, pH 8, 100 mM NaCl, 0.1 mM TCEP). The preparation of the GST-tagged AP2 complex has been described previously and followed the same steps as that of GST-tagged NTL-AP2 (32).

The preparation of the α-Nef/σ2 protein used for the FP assay in Figure 3 has been described previously (46).

### Binary binding test using size exclusion chromatography (SEC)

MBP-NTL and MBP-Nef (0.5 mg each) were mixed in a solution with a final volume of 500 μL and incubated on ice for 1 hr. The sample was then run through a Superdex 200 10/300 GL size exclusion column. MBP-NTL and MBP-Nef were also analyzed using SEC individually. The three elution profiles were overlaid to identify possible shifts. Other Serinc3 intracellular loop constructs were tested in the same fashion for possible binary binding with Nef.

### *In vitro* GST pulldown assay

Purified AP2^Δμ2-CTD^-GST or NTL-AP2^Δμ2-CTD^-GST (0.2 mg) and Nef (0.1 mg) were mixed in a final volume of 120 μl and incubated at 4 °C for 1 hr. The protein solution was then loaded onto a small gravity flow column containing 0.2 ml of GST resin. Flow-through was collected and the resin was extensively washed with 5 × 0.9 ml of the binding buffer (50 mM Tris, pH 8, 100 mM NaCl, 0.1 mM TCEP). The bound proteins were then eluted with 5 × 0.1 ml of GST elution buffer containing 10 mM reduced glutathione. The eluted proteins were analyzed by SDS–PAGE followed by Coomassie blue staining.

### Competition of TMR-cyclic-CD4_CD_ by unlabeled MBP-NTL/ICL peptides in the FP assay

For competition using unlabeled MBP-NTL, a stock solution of 120 µM unlabeled MBP-NTL was first prepared. The stock solution was then serial-diluted (2-fold each) 10 times. For making the final assay solutions, purified α-Nef/σ2 was buffer exchanged into the assay buffer (50 mM Tris, pH 8.0, 150 mM NaCl, 0.5 mM DTT, 0.01% Triton X-100). The α/σ2 protein (25 µM) and TMR-cyclic-CD4_CD_ (200nM) were first mixed and incubated at room temperature for 30 minutes. Then MBP-NTL was added at different concentrations, the plate was incubated at room temperature for 2 hours, and FP signals were then recorded. All experiments were done in triplicates. The FP data was fitted with non-linear regression using *OriginLab*. Statistical analysis was performed using Ordinary one-way ANOVA in *GraphPad Prism*. A p value of < 0.05 is considered statistically significant. Experiments using MBP-ICLs as competitors were performed the same way.

### Labeling NTL with the TMR fluorophore

NTL(Cys16)-encoding gene was cloned into a pMAT9 vector. Expression and purification of MBP-NTL(Cys16) was carried out the way as MBP-NTL described above. Conjugation of the TMR fluorophore onto NTL(Cys16) was done by following a published protocol (47). Briefly, 4 nmoles of purified MBP-NTL(Cys16) was buffer exchanged into TGED buffer (20 mM Tris, pH 7.9, 0.1 mM EDTA, 1 mM DTT, 5% Glycerol). The proteins were then fully reduced by adding DTT to 10 mM DTT, followed by 2 hr-incubation at 4 °C. Saturated ammonium sulfate solution was then added to the sample to precipitate the protein. The sample was then centrifuged at 13,000 rpm for 5 min at 4 °C and the supernatant was subsequently removed. The protein pellet was washed with buffer A (100 mM Na_2_PO_4_, pH 7.0, 200 mM NaCl, 1mM EDTA) several times *via* centrifugation. The protein pellet was then dissolved into 100 µl buffer A containing 20 nmoles tetramethylrhodamine-5-maleimide and incubated in room temperature for 30 minutes. The excess reagents were purified away through a desalting column, and labeled MBP-TMR-NTL was recovered afterwards. The MBP tag was subsequently removed through cleavage by the Mpro protease overnight, and the TMR-NTL was finally purified through a HiTrap Q anion exchange column.

### Fluorescence polarization assay using α-Nef/σ2 and TMR-NTL

The purified α(1–398)/σ2 (20 µM) and MBP-Nef (WT or mutant; 30µM) were first mixed and incubated at room temperature for 30 minutes. Assays were carried out in *Corning* 384-well black microplates (3820). In each well, 200 nM TMR-NTL peptide was mixed with the proteins in a total volume of 15 μl. Incubation was done for 1 or 2 hr at room temperature with minimal exposure to light. Fluorescence polarization (FP) was then measured using the *EnVision* plate reader (*Perkin Elmer*) with excitation at 535 nm and emission at 595 nm. Measured FP values in triplicates were averaged and subsequently plotted as a function of protein concentration in a logarithmic scale using *GraphD prism*.

### Plasmids and Mammalian Cells

Plasmid encoding pBJ5-Serinc3-HA was described previously (48). The C-terminal HA epitope tag was removed and an internal HA (YPYDVPDYA) placed at position 314 (extracellular loop 4) using overlap PCR and restriction digestion at NotI and EcoRI sites in the pBJ5 plasmid backbone. The proviral plasmids pNL4-3 and pNL4-3ΔNef have been described previously (49–51). The HEK293 cells were maintained in Dulbecco’s modified Eagle’s medium supplemented with 10% fetal bovine serum and penicillin/streptomycin.

### Serinc3 Virion Incorporation and Immunoblotting

To probe the contributions of residues to Nef-mediated antagonism Serinc3, incorporation of Serinc was measured in HIV-1 virions lacking Nef (ΔNef) produced from HEK293 cells co-transfected to express Serinc3 and Nef mutants. HEK293 cells seeded in 6-well plates were transfected with plasmid DNA encoding the HIV-1 molecular clone pNL4-3ΔNef (2.4 µg), pBJ5-Serinc3-HA (300 ng), pCINL Nef (500 ng) WT or indicated mutants, and pCINeo (empty plasmid) to total 4 µg DNA, using Transporter 5 transfection reagent (PolySciences). The following day, culture supernates were clarified of cellular debris by centrifugation (1000 x g), and the virions were partially purified from the supernates by ultracentrifugation of 1 mL at 23,500 x g through 20% sucrose cushions. Virion-producer cells were harvested and pelleted by centrifugation (300 x g). The viral and cell pellets were resuspended in Laemmli buffer containing 50 mM TCEP [Tris(2-carboxyethyl) phosphine; Sigma]. To avoid boiling and the consequent aggregation of Serinc3, the samples were sonicated (Diagenode Bioruptor) before protein separation on 10% denaturing SDS-PAGE gels, transfer onto PVDF membranes, and immunoblotting with the antibodies indicated below. Immunoreactive bands were detected using the Western Clarity detection reagent (Bio-Rad), and ChemiDoc imager system (Bio-Rad).

Primary and secondary antibodies were prepared in antibody dilution buffer, consisting of 1% milk in phosphate-buffered saline (PBS) with 0.02% Tween 20 (PBST). The following antibodies were used for detection of the proteins of interest: HA.11 (mouse; BioLegend), GAPDH (glyceraldehyde-3-phosphate dehydrogenase; mouse; GeneTex), HIV-1 p24 (mouse; Millipore), and HIV-1-Nef (Rabbit; Abcam). HRP-conjugated goat anti-mouse and donkey anti-rabbit secondary antibodies were from Bio-Rad.

## RESULTS

### Nef binds to the N-terminal cytoplasmic tail of Serinc3 *in Vitro*

High-resolution structures solved by us and others have shown that, when downregulating host membrane proteins, Nef typically binds to a short cytoplasmic segment of its target (32, 52, 53). Serinc3 is a multipass transmembrane protein and thus contains several intracellular loops: the N-terminal loop/tail (NTL), intracellular loop 1 (ICL1), ICL2, ICL3, ICL4, and C-terminal loop/tail (CTL) (Figure 2AB) (12). Two modes of binding are possible between Nef and Serinc3: 1) a single intracellular loop of Serinc3 is involved; 2) multiple intracellular loops are involved. In the related case of Nef-mediated Serinc5 downregulation, it has been shown that a single intracellular loop of Serinc5, namely ICL4, is mainly (if not solely) responsible. We hypothesized that Serinc3-Nef binding may similarly involve a single intracellular loop/segment of Serinc3. We also hypothesize that, even if more than one intracellular segment of Serinc3 are involved, a particular segment may be the main contributor and driver of the Serinc3-Nef interaction. To test our hypothesis, we decided to investigate each intracellular segment of Serinc3 individually for their possible binding with Nef. Among the Serinc3 intracellular segments, the CTL contains only six amino acids and is thus too short to be possibly targeted by Nef (Figure 2B). We therefore tested each of the remaining cytoplasmic loops of Serinc3—NTL, ICL1, ICL2, ICL3 and ICL4.

**Figure 2.**
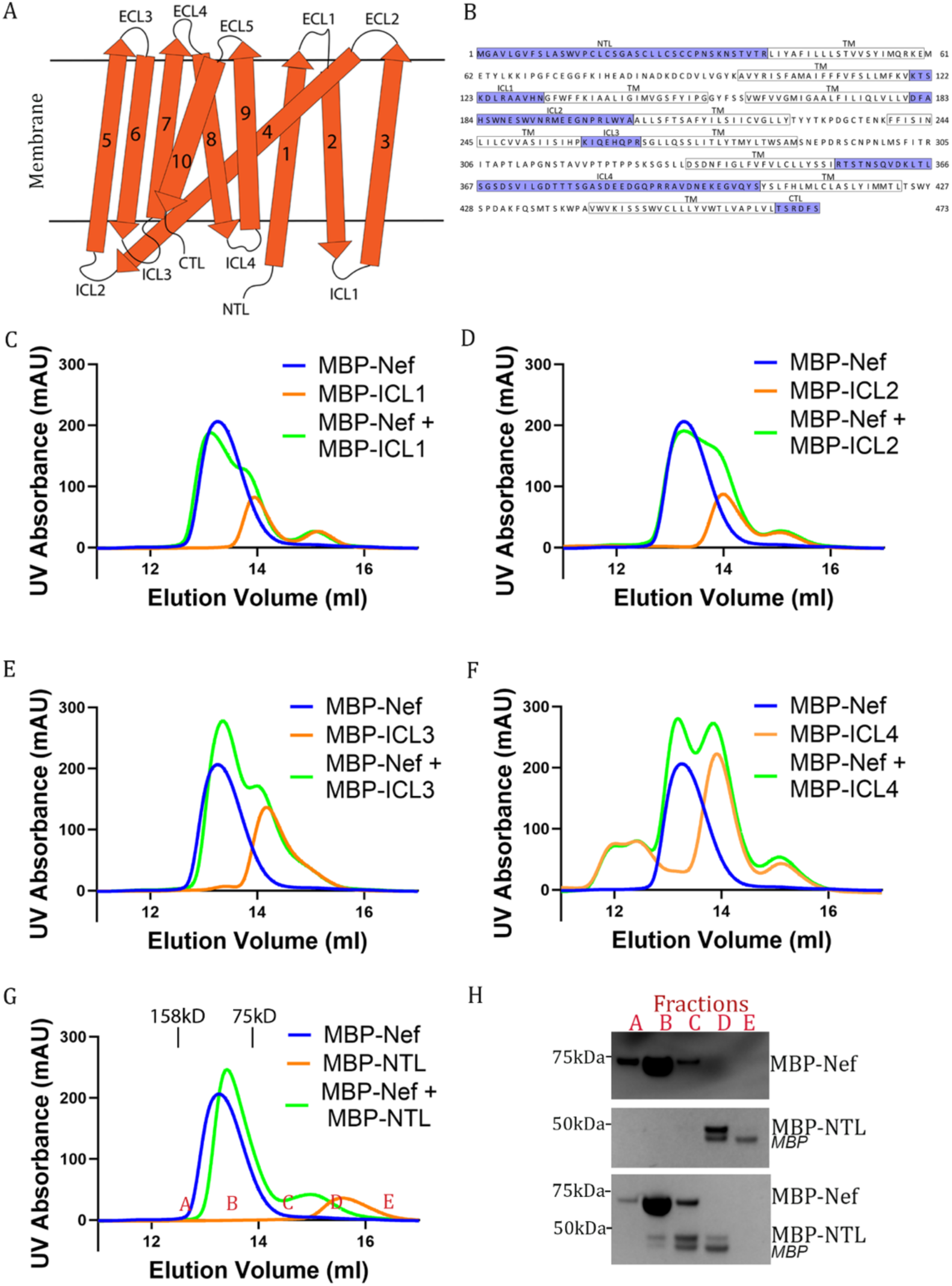
In vitro binding observed between Nef and the NTL of Serinc3, A) Schematic of Serinc3 topology as a transmembrane protein with intracellular (ICL) and extracellular loops (ECL). B) Amino acid sequence of Serinc3 with intracellualr segments highlited. C-F) SEC tests with MBP-ICLs showed that MBP-Nef does not bind to MBP-ICL1, MBP-ICL2, MBP-ICL3, or MBP-ICL4. G) In the SEC test, the mixture of MBP-Nef and MBP-NTL exhibited an elution profile (green curve) that differs from the combination of the MBP-Nef elution profile (blue curve) and the MBP-NTL elution profile (orange curve). H) SDS-PAGE analysis of elution fractions of SEC runs on MBP-Nef alone, MBP-NTL alone, and the mixture of the two.

Each Serinc3 loop was fused to the C-terminus of the maltose binding protein (MBP) and was subsequently expressed and purified. Nef was also expressed and purified as an MBP-Nef fusion. We then used a size exclusion chromatography (SEC)-based assay to detect possible binary binding between MBP-Nef and each of these MBP-Serinc3 loop fusions. When the Serinc3 ICLs were tested in the SEC assay, no shift was exhibited when the elution profile of the MBP-Nef:MBP-ICL mixture and the elution profiles of individual proteins are compared, indicating that Nef does not bind to any of the Serinc3 ICLs (Figure 2C-F). Intriguingly, however, when the Serinc3 NTL was tested, the elution profile of the mixture of MBP-NTL and MBP-Nef displayed a significant shift compared to the elution profiles of individual proteins (Figure 2G). The shift in proteins’ elution positions was confirmed by SDS PAGE analysis of eluted fractions (Figure 2H). These data show that there is binary interaction between Nef and the Serinc3 NTL.

We note that how the elution profiles of MBP-NTL and MBP-Nef shifted in the presence of each other was somewhat unexpected. In the presence of MBP-Nef, the elution of MBP-NTL shifted to the higher molecular weight (MW) region, which is consistent of complex-formation between MBP-NTL and MBP-Nef (Figure 2GH). On the other hand, however, in the presence of MBP-NTL, the elution of MBP-Nef shifted toward the lower MW region (Figure 2GH). On the first look, this shift of MBP-Nef seemed counterintuitive, because complex formation between two proteins typically results in both proteins being eluted in the higher MW region. However, careful analysis of this shift suggests to us that it may indicate an interesting aspect of the Nef-NTL binding. In the absence of MBP-NTL, MBP-Nef, which is 63.9 KDa in size, runs as a dimer on SEC (Figure 2G). Since MBP-NTL (47.2 KDa) is smaller in MW in comparison to MBP-Nef, a 1:1 complex of MBP-Nef and MBP-NTL should be smaller in MW than the dimer of MBP-Nef. Thus, the MBP-NTL-induced shift of MBP-Nef from higher to lower MW region is consistent with the scenario that MBP-Nef is in the monomeric form when binding to MBP-NTL. Importantly, a similar scenario had been observed in the Nef-CD4 association: the conformation of Nef required for CD4-binding is incompatible with Nef dimerization (32).

### Serinc3 NTL-binding and AP2-binding involve different parts of Nef and are compatible with each other

Nef-mediated Serinc3 downregulation, like Nef-mediated CD4 downregulation, involves hijacking of clathrin AP2-dependent endocytosis (61). The molecular details of the Nef-AP2 interaction—in the absence of CD4 or any other targeted host receptor/factor—was first elucidated structurally by Ren and colleagues in 2014 (31); their high-resolution crystal structure showed that Nef’s C-terminal loop forms an extensive interface with the α-σ2 hemicomplex of AP2. Subsequently, a high-resolution structure of Nef in complex with the hijacked tetrameric AP2 and the targeted CD4 cytoplasmic domain (CD4_CD_) was solved by us (32). In this structure, the Nef C-terminal loop makes the same exact interactions with the σ2 and α subunits of AP2 (Figure 1B). These structural findings argue strongly that the latching of the C-terminal loop of Nef onto AP2 is the foundational interaction and is very likely maintained in the Nef-mediated AP2-dependent downregulation of other host factors including Serinc3 and Serinc5.

The above reasoning led us to hypothesize that Nef, while engaged with AP2 through its C-terminal loop, may use a separate surface to bind and recruit Serinc3 NTL—a scenario similar to Nef-mediated AP2-dependent downregulation of CD4 (32). To test this, we designed and created a chimeric fusion protein between NTL and AP2. If our hypothesis is correct and the fusion design is appropriate, then the NTL-AP2 fusion should bind Nef at two separate sites— the NTL-binding site and the AP2-binding site—simultaneously and thus exhibit an affinity greater than that between AP2 and Nef.

In creating the chimeric NTL-AP2 construct, we fused the NTL of Serinc3 *via* a 10-amino-acid linker to the N-terminus of the β2 subunit of AP2. We chose this design because the very N-terminus of β2, according to our previous structure of CD4-Nef-AP2 (32), is located very close to the multifunctional pocket of Nef, which is frequently involved in binding and recruiting target proteins (32, 52). Thus, fusing NTL to the N-terminus of β2 should allow NTL to reach conveniently its potential binding site on Nef. The tetrameric complex containing the NTL-β2 chimera, the N-terminal domain of μ2 (μ2N), the σ2 subunit, and the α subunit fused with a C-terminal GST tag (α-GST) was co-expressed, and the assembled tetrameric complex, hereafter referred to as GST-tagged NTL-AP2, was subsequently purified. The purified protein suffered from some degradation of the NTL-β2 fusion but a significant portion of the protein nonetheless contained the complete NTL-β2 chimera (Figure 3, right lane of the loads).

**Figure 3.**
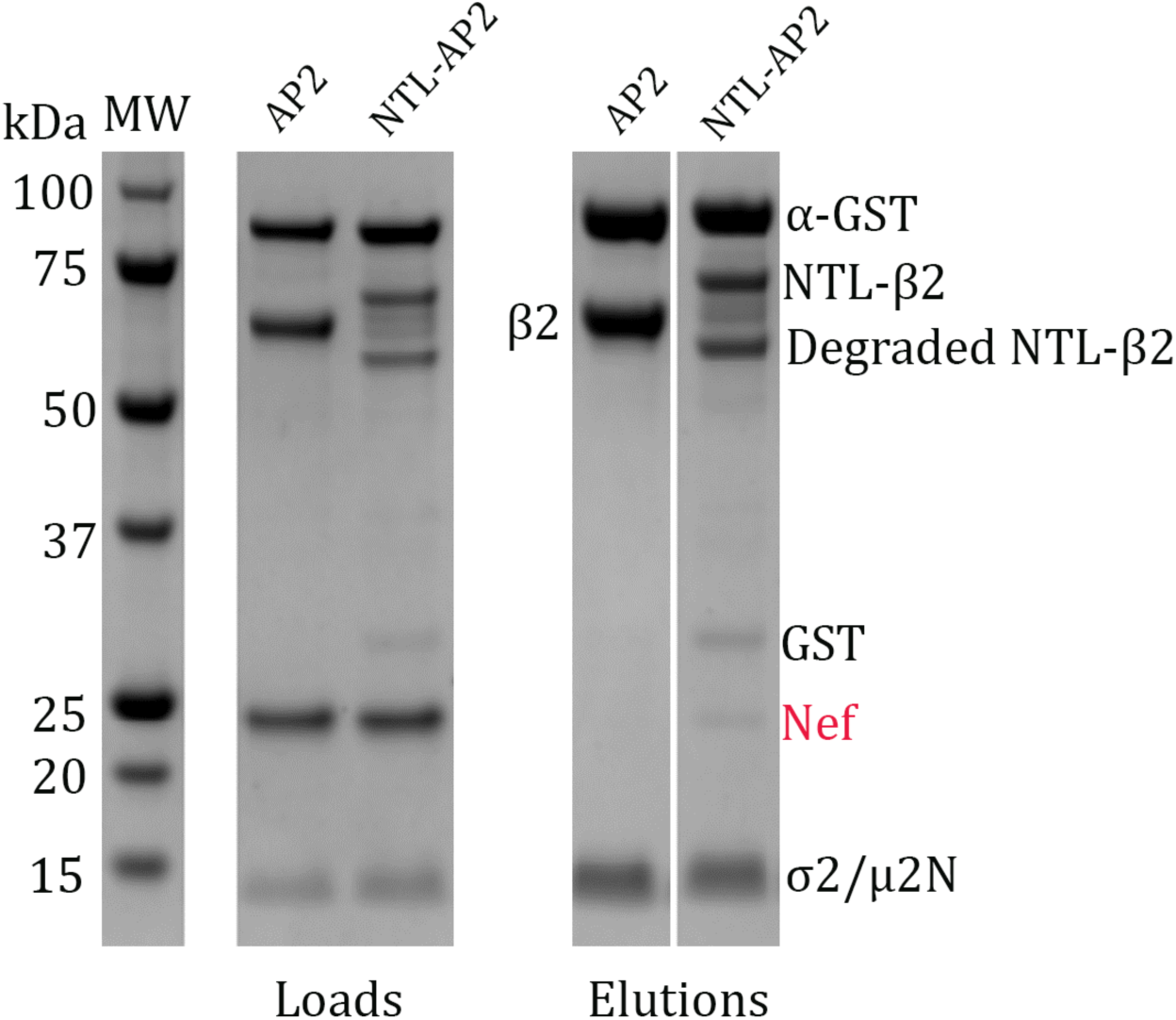
In vitro GST pulldown assay comparing NTL-AP2 and AP2 for binding with Nef. As shown in the SDS-PAGE of the loads (Coomassie blue staining), despite partial degradation in NTL-β2, purified GST-tagged NTL-AP2 protein was in the expected tetrameric state and exhibited a stoichiometry comparable to that of GST-tagged AP2. SDS PAGE analysis of the pulldown elution showed that, while no retainment of Nef was observed in the elution when GST-tagged AP2 was used as the bait protein, modest retainment of Nef was observed when GST-tagged NTL-AP2 fusion was used as the bait protein.

Using a GST-pulldown assay, we compared GST-tagged NTL-AP2 and GST-tagged AP2 for their abilities to bind Nef. Elution solutions of the pulldown were analyzed using SDS-PAGE followed by Coomassie blue staining. As shown in Figure 3, the GST-tagged NTL-AP2 protein modestly retained Nef, while GST-tagged AP2 did not retain Nef at all. It should be noted that this GST-pulldown, with Coomassie blue staining of the SDS-PAGE, is a stringent assay: weak interactions are often undetectable (such as the AP2-Nef interaction here, which is known and has been validated structurally (31, 32)) and only interactions with moderate-to-high affinities can be detected. Nonetheless, results of this pulldown assay support our hypothesis that NTL and AP2 bind to different surfaces of Nef. In addition, this data also corroborates the findings of the SEC binding assay (Figure 2GH), further supporting a direct binding between the Serinc3 NTL and Nef.

### Serinc3 NTL-binding and CD4_CD_-binding involve the same conserved pocket on Nef

As revealed by the *in vitro* binding assays above (Figure 2GH and 3), Nef and NTL interact directly with each other (Figure 2GH), and this interaction is compatible with the Nef-AP2 interaction (Figure 3). Such a scenario greatly resembles the CD4-Nef-AP2 interaction: Nef and CD4 cytoplasmic domain interact directly; CD4-binding and AP2-binding involve different parts of Nef and are compatible with each other (32). Such similarity suggested to us that the Serinc3 NTL might bind into the same conserved pocket of Nef, which has been shown to be responsible for binding and recruiting CD4 as well as for binding/recruiting MHC-I in Nef-mediated, clathrin AP1-dependent downregulation (32, 52). To test this, we used a previously established fluorescence polarization (FP) assay to investigate whether NTL and CD4_CD_ compete for binding to Nef (Figure 4A) (46). As described previously, in this assay a minimal construct, namely α-Nef/σ2, can associate with the fluorescent probe tetramethylrhodamine-labeled cyclic CD4_CD_ (TMR-cyclic-CD4_CD_) generating a significant FP signal (46). Importantly, this FP signal is sensitive to competitive binding taking place at the CD4-binding pocket of Nef; an unlabeled CD4 cytoplasmic tail peptide, when added, caused the decrease of the FP signal in a dose-dependent manner (46). When we added MBP-NTL to this assay, the FP signal decreased in a dose-dependent manner (Figure 4B), consistent with a direct competition between MBP-NTL and the CD4-mimetic fluorescent probe. In contrast, none of the MBP-ICLs caused any decrease of the FP signal (Figure 4B). These results strongly indicate that the NTL-binding occurs, at least partially, at the conserved CD4-binding pocket of Nef. They also further confirm that Nef specifically recognizes and binds the Serinc3 NTL but not any of its ICLs.

**Figure 4.**
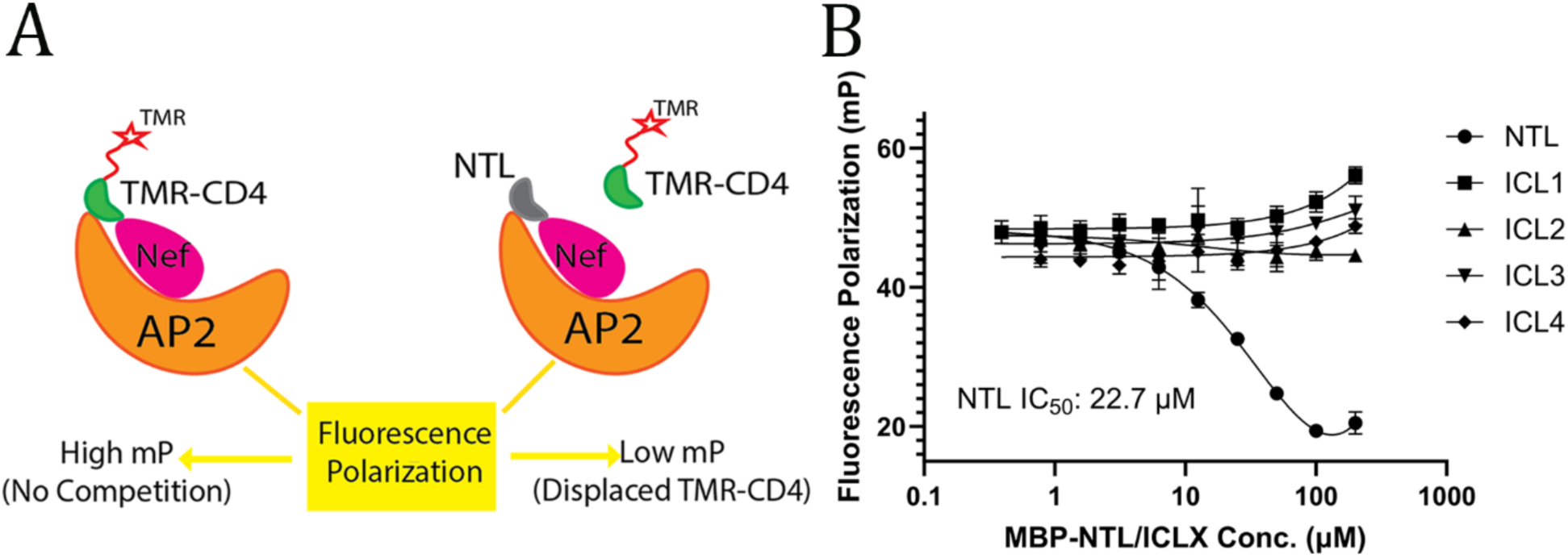
Serinc3 NTL and the CD4 cytoplasmic domain bind to the same multifunctional pocket of Nef. A) Cartoon illustrating the design of the fluorescence polarization assay. B) Addition of MBP-NTL led to a dose-dependent decrease of the FP signal (IC_50_: 22.7 µM) while no signal reduction was observed when any MBP-ICL was added. Analysis using one-way ANOVA indicated that the data of MBP-NTL competition are statistically significant (p = 0.0006).

### CD4_CD_- and Serinc3 NTL-binding share same determinants in Nef and should involve the same conformation of Nef

Our results discussed above pointed to some significant similarities between Serinc3 NTL and CD4 cytoplasmic domain in binding Nef and AP2. Encouraged by these findings, we then tested, through *in vitro* mutagenesis, whether Nef residues important for CD4-binding are also required for binding Serinc3 NTL. Here, a modified FP assay was used to monitor the binding between NTL and the Nef-AP2 complex: a TMR-labeled NTL peptide was used as the fluorescent probe. When this probe was introduced to the complex containing wild-type Nef and the α-σ2 hemicomplex of AP2, a significant FP signal was observed compared to the background (Figure 5, WT *vs*. control), which is consistent with an association taking place between TMR-NTL and the Nef-containing complex.

**Figure 5.**
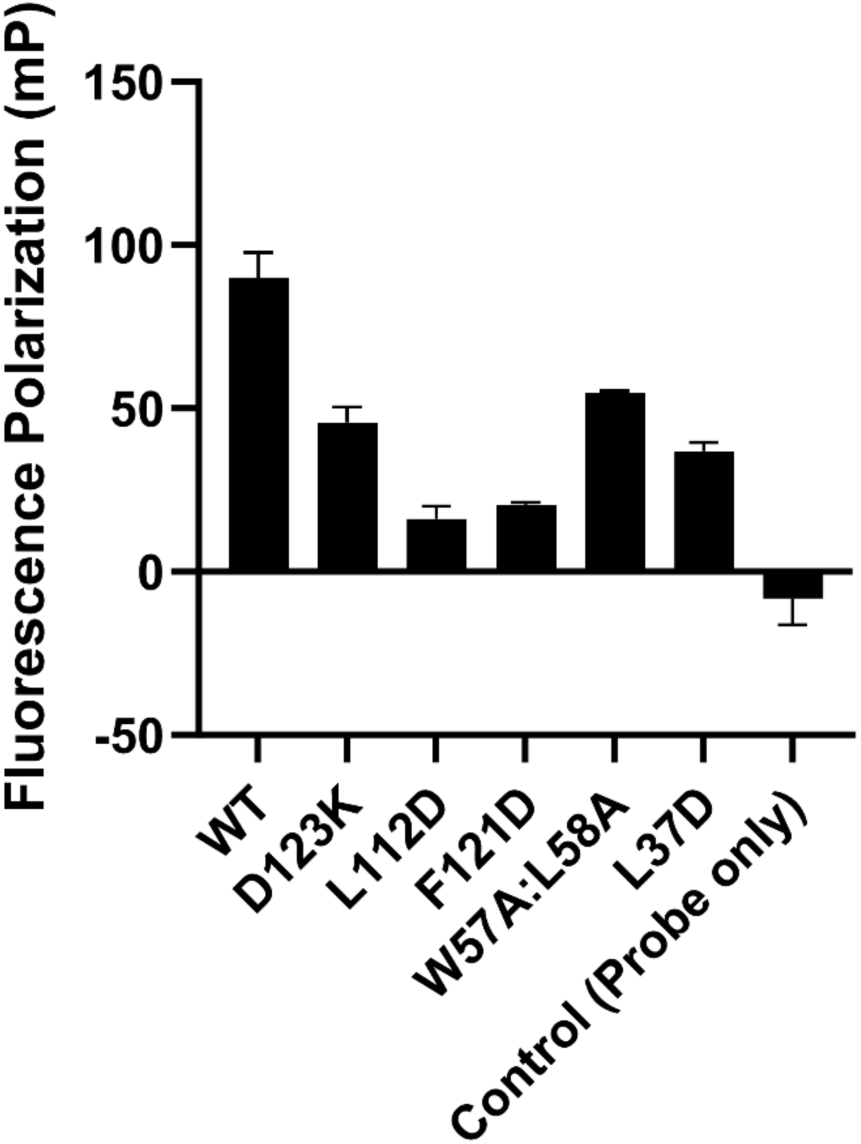
In vitro mutagenesis study revealed Nef residues important for Serinc3 NTL-binding. In the presence of α(1–398)/σ2 and the TMR-NTL probe, wild type (WT) Nef produced a FP signal of ∼90 mP in comparison to the control (probe only). All Nef mutations tested led to a reduction of FP signal to different degrees. Data are shown as mean and standard deviation of three technical replicates.

We then investigated whether Nef mutations known to disrupt Nef-CD4 binding could similarly disrupt the Nef-NTL association. The specific mutations tested are D123K, L112D, F121D, W57A:L58A, and L37D. Notably, not all Nef residues tested here contribute to CD4-binding through direct contact with CD4. While Nef residues Leu37, Phe121, and Asp123 make direct contacts with CD4, Trp57, Leu58, and Leu112 do not (32) (Figure 1C). To enable CD4 binding and downregulation, Trp57 and Leu58, which are located on a short helix within the Nef N-terminal loop, dock into a hydrophobic pocket on Nef core formed by Leu112, Phe121 and other residues (Figure 1C); this intramolecular association stabilizes a specific conformation of the Nef N-terminal loop and positions Leu37 and neighboring residues to bind CD4 (32).

Remarkably, when tested in our FP-based binding assay, each of the Nef mutants exhibited decreased FP signals indicating that the binding to TMR-NTL was compromised by the mutation (Figure 5). These results revealed that these Nef residues (Asp123, Leu112, Phe121, Trp57, Leu58, and Leu37), which had been shown to be required in CD4 binding and downregulation by Nef, are also involved in the Nef-NTL association. Moreover, our results here suggest that the unique Nef conformation, which is critical for CD4-binding and downregulation (Figure 1C), should be similarly required for the Nef-NTL binding.

### Virion exclusion of Serinc3 requires the same Nef residues important for CD4 downregulation

As shown above, results from our *in vitro* tests converge and collectively suggest that Nef binds to the NTL of Serinc3 and likely downregulates Serinc3 through a mechanism similar to that of CD4 downregulation. We then tried to validate these findings in human cells. Here, we used a previously established assay to measure Nef-mediated exclusion of Serinc3 from virions by western blot (48). Nef-negative (ΔNef) HIV-1 virions were produced from HEK293 cells in the presence of Serinc3 containing an HA epitope tag, either with or without co-expression of wild type Nef or Nef mutants. As expected, Serinc3 was detected in partially purified virions (Figure 6). The expression of wild-type Nef reduced the amount of Serinc3 in virions (compare to the Nef-negative control). In contrast, expression of Nef encoding the L164A:L165A mutation, which is known to disrupt the Nef-AP2 interaction (31, 32), restored the level of Serinc3 in virions relative to wild type Nef. In a similar fashion, mutations of residues within the CD4-binding pocket, F121D and D123R, which are known to disrupt CD4 binding and downregulation (32–34), also restored the levels of Serinc3 in virions, respectively (Figure 6). These data confirm our observations *in vitro* and further prove that the conserved pocket of Nef is indeed involved in recruiting Serinc3 in cells. We also tested the L112D and the W57A:L58A mutations, which as discussed above should prevent Nef from adopting the distinct conformation critical for CD4 binding and downregulation. Consistent with our in vitro mutagenesis results, these mutations also inhibited Nef-mediated exclusion of Serinc3 from virions (Figure 6).

**Figure 6.**
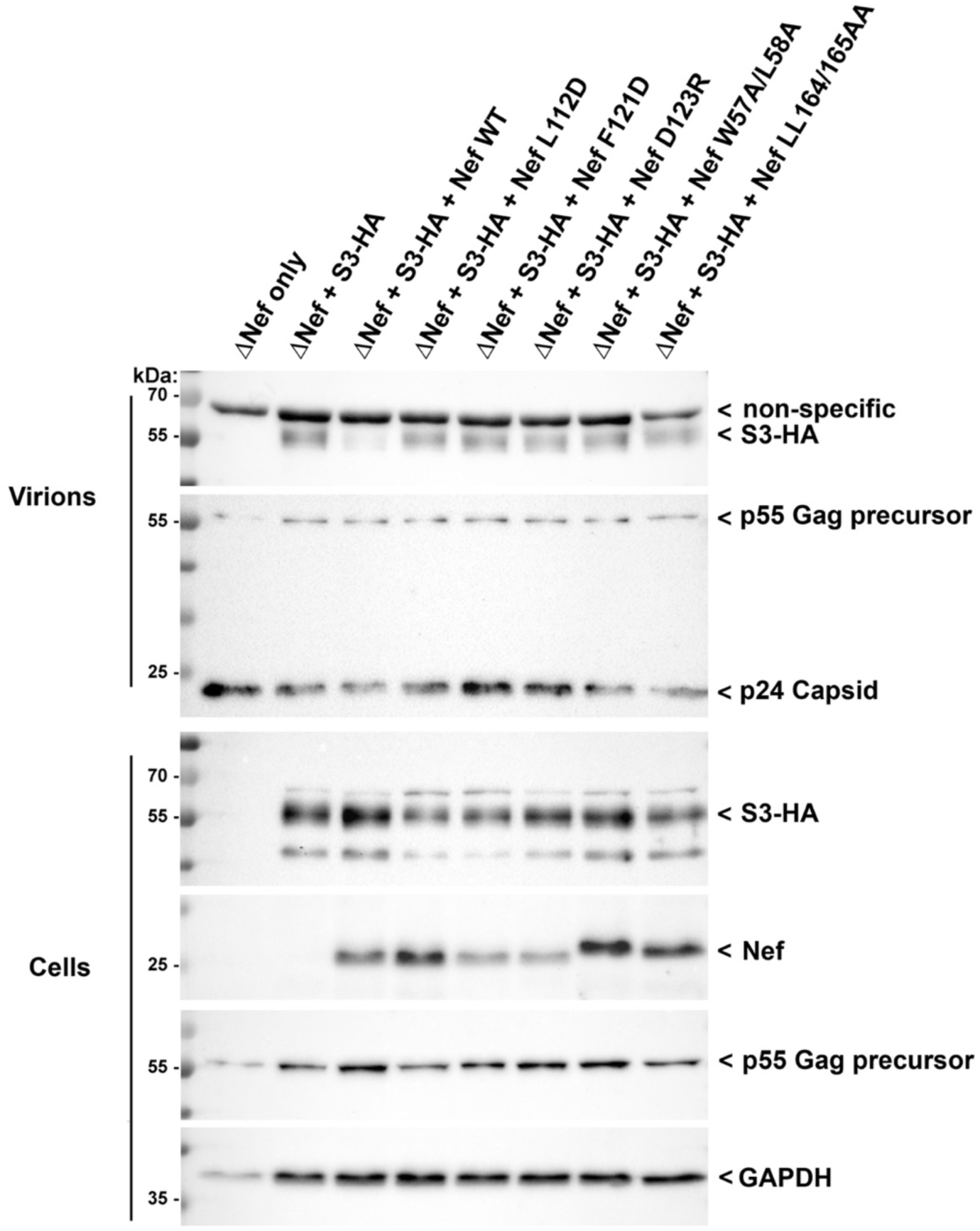
Nef residues important for CD4 downregulation are also required for excluding Serinc3 from virions. Nef-negative HIV-1 virions were produced in HEK293 cells co-transfected to express HIV-1 lacking an intact nef gene (ΔNef), Serinc3 (HA-tagged; S3-HA) and wild type Nef (Nef WT) or the indicated Nef mutants, each from separate plasmids. Serinc3 (HA) was detected in virions, which were partially-purified by removal of cellular debris followed by centrifugal pelleting through a 20% sucrose cushion as described in Methods, and in virion producer cell lysates by SDS PAGE and western blot. The blots were also probed for Nef, GAPDH (a cellular protein used as laoding control), and p24/p55 Gag. p24 is the virion-capsid antigen. p55 Gag precursor is the predominant form of Gag in cells, while in virions p24 predominates due to intra-virion processing of p55 by the viral protease. A non-specific band detected by the anti-HA antibody is indicated; this was observed in the virion preparations even in the absence of S3-HA expression (far left sample lane; top blot).

## DISCUSSION

Previous work on Serinc5 showed that HIV-1 Nef targets the ICL4 of Serinc5 for downregulation and antagonism (37). It has been subsequently speculated that Serinc3 downregulation may similarly involve Nef binding to the ICL4 of Serinc3 (12). However, results from our *in vitro* assays—a SEC binding assay, a GST pulldown assay, and two FP-based assays—converged in supporting that Nef binds directly to the N-terminal cytoplasmic tail of Serinc3 (Figures 2-5) but not any of its ICLs (Figures 2&4). A limitation of our study is that we tested each intracellular segment of Serinc3 separately, which should have prevented us from evaluating whether multiple loops of Serinc3 could be involved in Nef/AP2-binding simultaneously. We believe, however, that, even if multiple loops of Serinc3 are involved, the NTL of Serinc3 should still be a major contributor to the Nef-Serinc3 association. In support of this, we showed that mutations of Nef residues, which disrupted Nef-NTL association (Figure 5), also abolished Nef’s activity in excluding Serinc3 from budding virions (Figure 6).

Although Serinc5 and Serinc3 are similar both structurally and functionally (3, 4, 12, 54), different segments of Serinc5 and Serinc3 are targeted by Nef for their respective downregulation: Nef antagonizes Serinc5 through binding to its ICL4, while the downregulation of Serinc3 involves Nef associating with the N-terminal tail of this host factor. This finding highlights the functional versatility of Nef. It also suggests that, despite that Serinc3 is less potent than Serinc5 in restricting HIV-1 (3, 4), Serinc3 modulation is unlikely a “side-effect” of Serinc5 antagonism but is more likely a deliberate function of Nef shaped through evolution. A recent study revealed that Serinc3 is present in extracellular vesicles (EVs) and that its distribution within different populations of EVs is altered upon HIV-1 infection in a Nef-dependent manner (55). Conceivably, the Nef-mediated incorporation of Serinc3 into EVs, which is unique to Serinc3 and not observed in Nef’s modulation of other cellular targets (55, 56), may benefit HIV-1 in way(s) beyond merely restoring viral infectivity. It also suggests that, in addition to the plasma membrane, Nef may interact with Serinc3 in other cellular compartment(s) (e.g., multivesicular bodies, a place known for the biogenesis of EVs) to modulates its transportation and/or cellular level.

Surface downregulation of the Serinc proteins, like that of CD4, occurs through Nef hijacking of clathrin AP2-mediated endocytosis (4, 28, 29). As revealed by our previous crystal structure of Nef in complex with CD4_CD_ and AP2, while Nef uses mainly its C-terminal loop to bind AP2, CD4-binding involves a separate set of Nef residues and requires Nef adopting a specific conformation (32). Our work here shows that the same binding pocket as well as the same conformation should be involved in Serinc3 modulation: mutation known to disrupts the CD4-Nef interaction (D123K), mutations known to prevent Nef from adopting the specific conformation for CD4 binding (W57A:L58A, L112D), and mutation affecting both (F121D) all disrupted Nef-Serinc3 NTL binding (Figure 5) as well as Nef-mediated virion incorporation of Serinc3 (Figure 6) (32).

The downregulation of Serinc5 also shares many of the same determinants in Nef. Mutations involving Trp57, Leu112, and Phe121 in Nef each impaired Serinc5 downregulation (11, 38); the same effect was also observed when the Nef N-terminal residues 12-39 was truncated (40). While these and other findings point to great similarities between the modulation of Serinc5, Serinc3, and CD4, differences between the determinants of these Nef functions have also been reported. Studies, including those on Nef polymorphisms, showed that Serinc5 downregulation and CD4 downregulation are to some extent genetically separable—some Nef mutations differentially affect and thus potentially uncouple these two functions (42–45). Similarly, Nef proteins from different subtypes of HIV-1 (57), as well as from different patients (58), vary in their abilities in antagonizing Serinc5 and Serinc3, suggesting that these might also be genetically separable functions of Nef. Complete elucidation of the mechanisms of Nef-mediated downregulation of Serinc3/5 will require high-resolution structures of the corresponding complexes; we aim to solve such structures in the future.

Finally, the revelation that Serinc3-binding occurs at the conserved Nef pocket—the same site involved in recruiting CD4 and MHC-I (32, 52)—further underscores the significance of this pocket in drug discovery against Nef. Small molecules that bind avidly to this site should be able to rescue several cellular targets of Nef at once, which may revitalize multiple adaptive and innate immune mechanisms to combat HIV-1 infection. Such therapeutic(s), if developed successfully, could be of great impact as they are directly applicable to HIV-1 cure strategies.

## Data Availability

all data are contained within the manuscript.

## Author Contributions

M.K.S., C.S., J.G., and X.J. designed the experiments. M.K.S. performed the *in vitro* biochemical studies. C.S. and A.D. carried out the cellular studies. T.F. contributed to the SEC assays. All authors analyzed the data. X.J. conceived the project. X.J. and J.G. supervised the project. M.K.S., J.G., and X.J. wrote the manuscript.

## Acknowledgements

We thank Madelyn Davis, Priya Sridharan, and Fatema Yeasmin for their technical assistance. This work was supported by National Institutes of Health grants R01AI176897 (X.J.) and R01AI129706 (J.G.). The content is solely the official views of the authors and does not necessarily represent the official views of the National Institutes of Health. This publication was supported in part by the James B. Pendleton Charitable Trust and by the San Diego Center for AIDS Research (SD CFAR), an NIH-funded program (P30 AI036214), which is supported by the following NIH Institutes and Centers: NIAID, NCI, NHLBI, NIA, NICHD, NIDA, NIDCR, NIDDK, NIGMS, NIMH, NIMHD, FIC, and OAR.

## Conflict of Interest

the authors declare that they have no conflicts of interests with the contents of this article.

## Notes

### Competing Interest Statement

The authors have declared no competing interest.

